## Extended data figures for "An efflux pump family distributed across plant commensal bacteria conditions host- and organ-specific detoxification of a host-specific glucosinolate"

**Extended data Figure 1. The abundance of Paraburkholderia in wild soil**

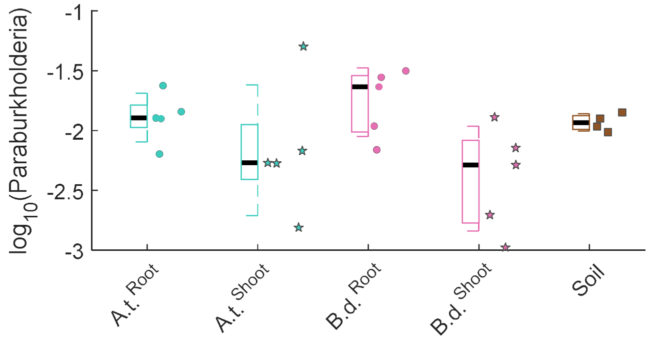


The Log transformed relative abundance of all members of the Paraburkholderia genus in the roots and shoots of both *Arabidopsis thaliana* Col-0 (A.t.) and *Brachypodium distachyon* BD-21(B.d.) as well as in the soil as calculated based on read counts of 16S rRNA gene amplicon sequencing. Paraburkholderia are detectable and maintain similar levels in all sample types and are stable across most samples.

**Extended data Figure 2. Number of barcodes per sample in colonization assay**

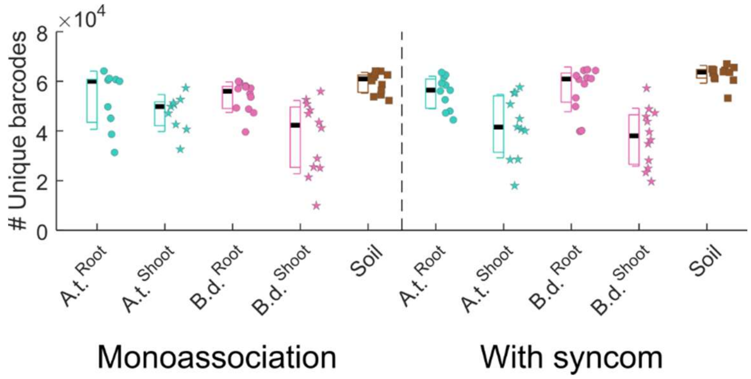


The number of unique TnBarSeq barcodes identified for each biological sample. Boxplots show the median (black line), 1$^{st}$ and 3$^{rd}$ quartiles (box), and one standard deviation (whiskers) of each sample. Different sample types show similar numbers of unique barcodes. Adding the TnBarSeq library alone (Monoassociation, left) or with a small auxiliary community of four members (With syncom, right) had no effect on the number of unique barcodes. As we recovered barcodes from 5,962 MF376 genes and the average number of unique barcodes per sample is 50,544; each sample had an average of ~8.5 unique barcodes per gene.

**Extended data Figure 3. Volcano plots of *Paraburkholderia bryophila* MF376 mutants in all habitats.**

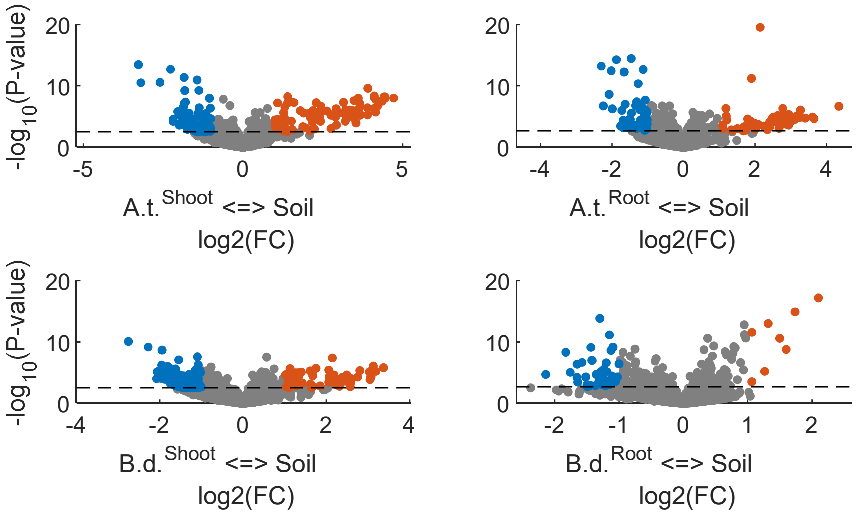


Volcano plot comparing the abundances of MF376 mutants in the soil to their abundances in association with different plant tissues. Fold changes (FC) and P-values for individual mutants were calculated based on soil samples and plant samples. Differentially abundant mutants that were statistically different (significance threshold, FDR corrected P-value < 0.05) are either depleted in association with plant tissue (plant association, blue, Log2(FC) < -1) or enriched in association with plant tissue (negative plant association, red, Log2(FC) > 1). All other mutants are defined as neutral (gray).

**Extended data figure 4. TnBarSeq results from *Pseudomonas simiae* WCS417 recapitulates our findings using *Paraburkholderia bryophila* MF376**

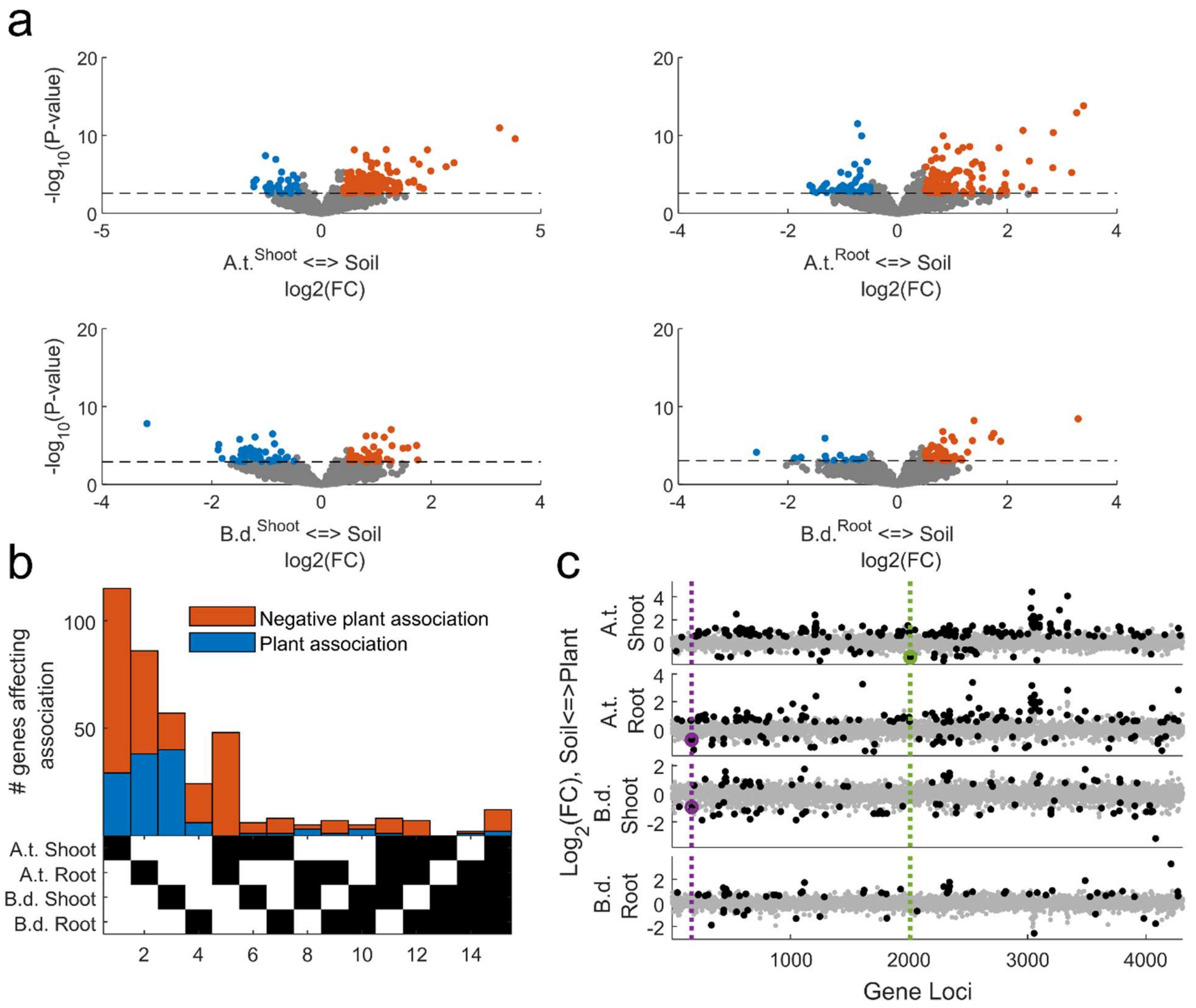


**a,** Volcano plot comparing the abundances of WCS417 mutants in the soil to their abundances in association with different plant tissue. Fold changes (FC) and P-values for individual mutants were calculated based on soil samples and plant samples. Differentially abundant mutants that were statistically different (significance threshold, FDR corrected P-value < 0.05) are either depleted in association with plant tissue (plant association, blue, Log2(FC) < -0.5) or enriched in association with plant tissue (negative plant association, red, Log2(FC) > 0.5). All other mutants are defined as neutral (gray). As the recovery of WCS417 barcodes was not as successful as that of MF376 and only few genes satisfied the threshold of Log2(FC) > |1|, we reduced the stringency of our criteria to Log2(FC) > |0.5|. **b,** An upset plot of genes that affect plant association by WCS417. Genes that positively (blue) or negatively (red) affect association with different host and organ combinations (bottom, black squares). Full list of genes is in extended data table 3. Like Fig 2a (for MF376) most WCS417 plant association genes are highly specific to both host and organ. **c,** Locus position in the WCS417 genome of statistically significant mutants in plant association (black dots). Organ specific efflux pump systems benefit the association with Arabidopsis shoots (green) or roots (purple) but not with other organs and host. Log2(FC) of mutant abundance in association with plant tissue compared to soil is plotted against their location on the genome. A single subunit of the root specific pump (a periplasmic gene) is also identified as a plant association gene in *Brachypodium distachyon* BD21 shoots.

**Extended data figure 5. Prevalence of *ef90* orthologs across bacteria from different habitats.**

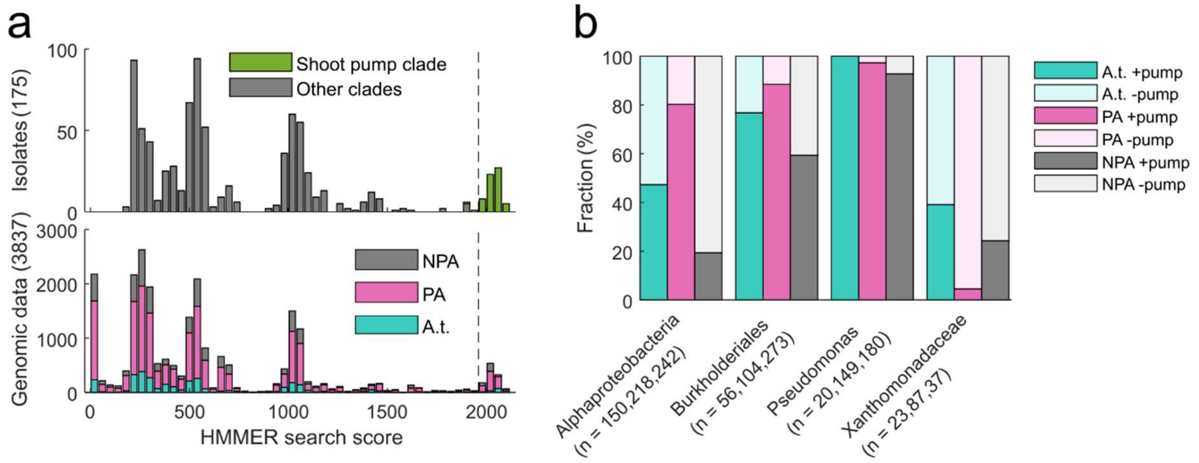


**a,** *ef90* homologs in the shoot-association clade are prevalent among bacteria from different habitats. A Hidden Markov Model (hmm) profile was built from members of the ef90 orthologous group using HMMER. (top) A histogram of the HHMER search score for the shoot clade hmm profile in the 175-member isolate collection derived largely from *Arabidopsis thaliana*. A strict HMMER threshold score (dashed line) includes the vast majority of genes in the shoot-association clade (*ef90 pumps*, green) and no other efflux pump genes (gray). (bottom) The HMMER search scores among genes in a large genomic database of 3837 bacterial genomes of Arabidopsis-associated (A.t., turquoise), non-Arabidopsis-plant-associated (PA, magenta), and non-plant associated (NPA, gray) bacteria. Genes in the shoot clade are right of the dashed line according to the HMMER threshold score found in the top panel. **b,** The prevalence of genomes that have a gene in the shoot-association clade in four Pseudomonadota taxonomic groups. Genomes that have at least one gene in the shoot-association clade of efflux pumps (+*ef90* pumps, dark color) are more prevalent among Arabidopsis-associated bacteria (A.t., turquoise) than among non-plant associated (NPA, gray) bacteria in all four taxonomic groups tested. Compared to no-Arabidopsis-plant-associated bacteria (PA, magenta), *ef90* orthologs are either more (Pseudomonas and Xanthomonadaceae) or less (Alphaproteobacteria and Bukholderiales) prevalent among Arabidopsis-associated bacteria.

**Extended data figure 6. The *saxF* efflux system from *Pseudomonas syringae* pv. tomato DC3000 is a member of the *ef90* orthologous group.**

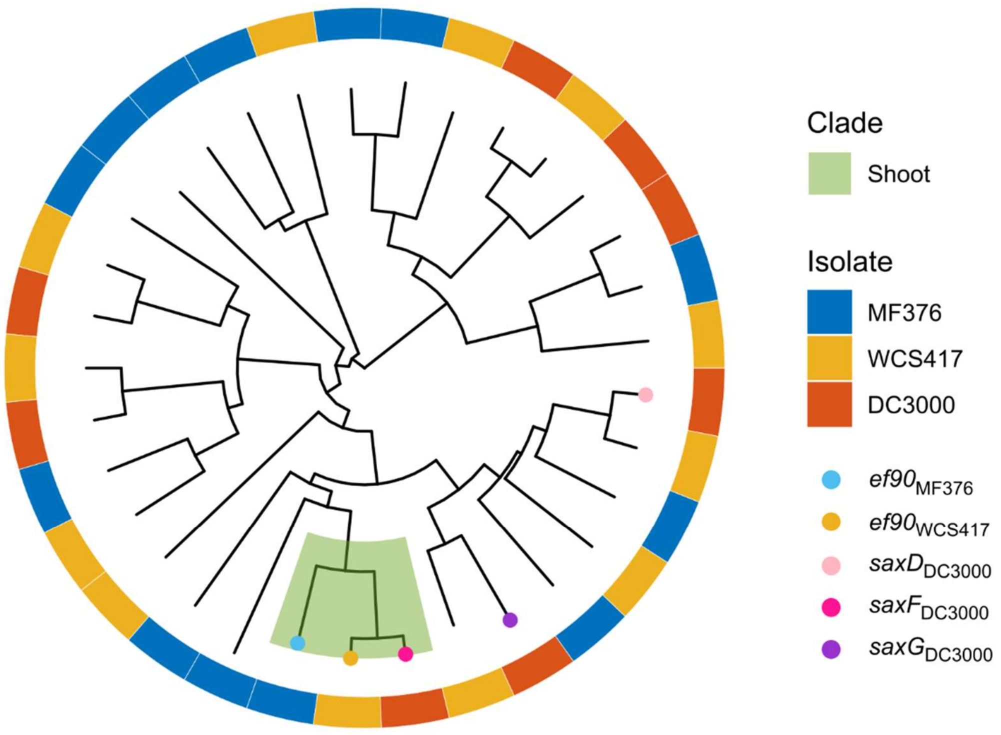


A gene tree of all inner membrane subunit RND-type efflux pumps from *Paraburkholderia bryophila* MF376, *Pseudomonas simiae* WCS417, and *Pseudomonas syringae* pv. *tomato* DC3000. The *saxF* gene from DC3000 sits within the shoot association clade defined by the MRCA of the ef90 orthologs from MF376 and WCS417.

**Extended data figure 7. The *ef90* ortholog in *Pseudomonas simiae* WCS417 does not confer resistance to shoot extract of either *Capsella bursa* or *Arabidopsis thaliana* Ler-1**

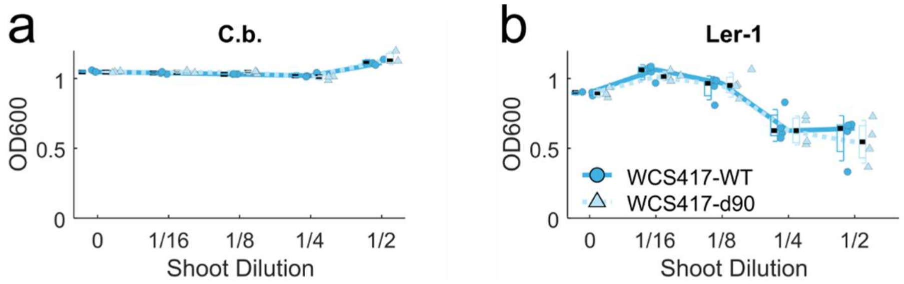


Fitness of *Pseudomonas simiae* WC417 d90 in macerated leaf extract. Final OD of overnight culture of wildtype *Pseudomonas simiae* WCS417 (WCS417-WT, dark cyan, solid line) or the mutant in the homologous shoot association efflux pump (WCS417-d90, light cyan, dashed line) in medium supplemented with diluted macerated leaf extract from *Capsella bursa* (**a**) or *Arabidopsis thaliana* Ler-1 (**b**). The mutant's growth inhibition seen for *Arabidopsis thaliana* Col-0 (Fig. 3d) is not seen for either host.

**Extended data figure 8. Expression *ef90/saxF* is induced by shoot extract from *Arabidopsis thaliana* Col-0 but not by *Capsella bursa-pastoris***

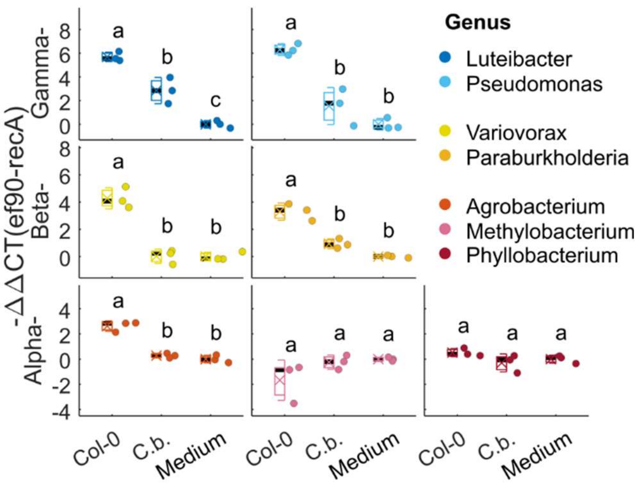


The expression of the ef90 pump of representative strains was measured using RT-qPCR 15 minutes after exposure to stimulus. The ef90 pump is induced to higher levels than by C.b. by five of seven representative strains (Fig. 5e) from all classes (Anova with post-hoc Tukey HDS test, P-value<0.05)

**Extended data figure 9. The diverse bacterial inhibitory profile of Arabidopsis thaliana genotypes**

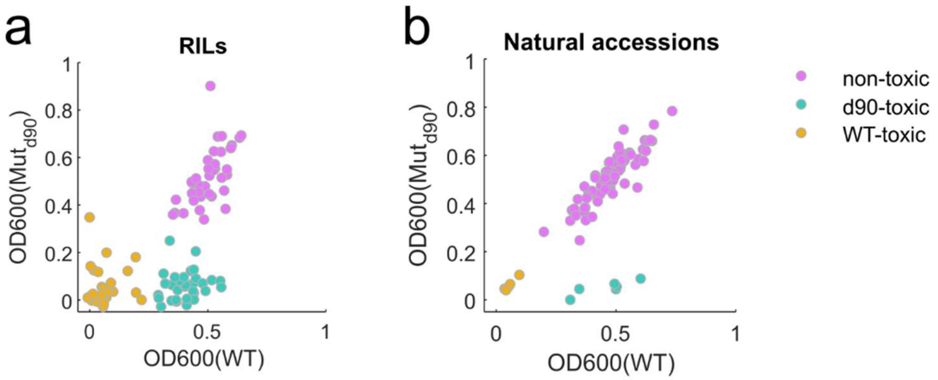


Final optical density of wildtype (WT) and the mutant d90 of Paraburkholderia bryophila MF376 after an overnight growth with leaf extract of Col-0xLer-0 RILs (**a**) or natural accessions (**b**). In both populations we identify three groups with distinct inhibition profile: non-toxic, don't inhibit bacterial growth (magenta); d90-toxic, inhibit only the growth of the d90 mutant and not of the WT bacteria (turquoise); WT-toxic, inhibit the growth of both mutant and WT bacteria (yellow).

**Extended data figure 10. The three phenotypic groups diverge in ESP expression and dominant glucosinolate chemotype**

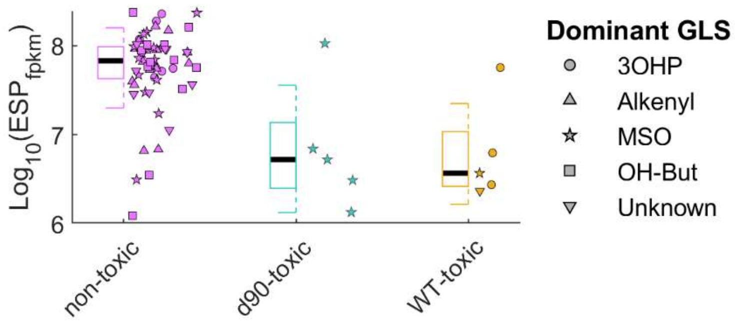


The expression level of ESP and the dominant chemotype of natural accession divided by phenotypic group. The expression of ESP is higher among non-toxic accession compared to both toxic groups. Looking at the dominant glucosinolate type we show that all d90-toxic accessions have MSO-GLS and three of the known four WT-toxic accessions have 3OHP-GLS, both in agreement with Fig. 6c. On the other hand, only one of the six non-toxic accessions that express low levels of ESP have either of them (the one outlier is accession 7378, Uk-1, with 4MSO-GLS).

**Supplementary tables**

Supplementary table 1. Plant association genes for *Paraburkholderia bryophila* MF376

Supplementary table 2. Enriched COG categories

Supplementary table 3. Plant association genes for *Pseudomonas simiae* WCS417

Supplementary table 4. Members of the ef90 orthologous group

Supplementary table 5. Representative strains used for RT-qPCR validation

Supplementary table 6. Phenotypic and genotypic profiles of Col-0 x Ler-1 RILs

Supplementary table 7. Phenotypic profiles of natural *Arabidopsis thaliana* accessions

Supplementary table 8. GWA results

Supplementary table 9. Primers list
